# SCM-1/SCAMP Maintains Microdomain Boundaries and Cargo Sorting within the Endosomal System

**DOI:** 10.64898/2026.05.20.726532

**Authors:** Kam Shan Hu, Anne Norris, Wilmer Rodriguez-Polanco, Collin McManus, Inna A. Nikoronova, Geoffrey G. Hesketh, Anne-Claude Gingras, Maureen M. Barr, Barth D. Grant

**Affiliations:** Department of Molecular Biology and Biochemistry, Rutgers University, Piscataway, NJ 08854; New York Structural Biology Center, The City College of New York, New York 10027, USA; Department of Genetics and Human Genetics Institute of New Jersey, Rutgers, The State University of New Jersey; Piscataway, New Jersey 08854, USA; Center for Lipid Research, New Brunswick, NJ, USA; Lunenfeld-Tanenbaum Research Institute, Mount Sinai Hospital, Sinai Health, Toronto, Canada; Department of Molecular Genetics, University of Toronto, Toronto, Canada

## Abstract

After endocytosis, transmembrane cargo reaches sorting endosomes where it is partitioned into physically distinct recycling or degradative microdomains. While the J-domain protein RME-8/DNAJC13 is known to maintain these boundaries by actively removing degradative machinery from the recycling microdomain, other factors that contribute to this spatial organization remain poorly defined. Here, we identify the conserved tetraspan protein SCM-1/SCAMP as a key microdomain organizer, discovered through RME-8 proximity-dependent biotinylation screens in *C. elegans* and human cells. Leveraging the large endosomes of *C. elegans* coelomocytes, we show that SCM-1 is selectively enriched within the recycling microdomain. In *scm-1* mutants, recycling and degradative microdomains still assemble but fail to remain spatially distinct, resulting in inappropriate microdomain overlap. This loss of boundary integrity occurs without increasing the recruitment of sorting machineries, indicating a mechanism distinct from the RME-8-mediated uncoating pathway. *scm-1* mutants exhibit significant sorting defects, including misrouting of recycling cargo MIG-14/Wls and v-SNARE SNB-2/VAMP3 to late endosomes and lysosomes. We find that *snb-2* mutants themselves missort MIG-14 to late endosomes and lysosomes, suggesting that SNB-2 sorting is key for recycling function. Our data suggest that both microdomains lose efficiency in *scm-1* mutants, as cargo missorted into late endosomes and lysosomes is not depleted overall, and degradation of an independent ESCRT-dependent cargo is delayed. We conclude that SCM-1 ensures endosomal sorting fidelity by stabilizing microdomain boundary integrity, a process required for efficient recycling and degradation of transmembrane cargo.

## Introduction

After endocytosis, a complex mixture of internalized plasma membrane cargo enters endosomes, processing centers for sorting of endosomal content. Peripheral membrane coat complexes of the endosome recognize and sort cargo, directing subsets toward lysosomal degradation or recycling pathways leading to the Golgi (retrograde recycling) or the plasma membrane, either directly or via recycling endosomes.^1–4^

Increasing evidence indicates that the competing sorting activities of the endosome are organized into physically distinct microdomains on the endosomal limiting membrane. Retromer and associated sorting nexins occupy recycling microdomains that promote cargo retrieval through tubular carriers, whereas ESCRT machinery occupies neighboring degradative microdomains that recognize ubiquitylated cargo and drive such cargo into intraluminal vesicles (ILVs) that are ultimately degraded in the lysosome.^5–11^ Physical separation of these competing microdomains has been proposed to increase sorting efficiency and prevent interference between opposing trafficking pathways.^12^

Our previous work identified the conserved J-domain protein RME-8 as a key regulator of microdomain separation.^8^ Studies in the *C. elegans* intestine were the first to show that RME-8 binds directly to sorting nexin SNX-1 and participates in endosome-to-Golgi recycling, properties that are conserved in mammals.^13–15^ Further analysis showed that RME-8 and SNX-1 control endosomal clathrin dynamics, which was initially surprising because endosomal clathrin is thought to form a flat lattice associated with the degradative ESCRT machinery within degradative microdomains rather than acting within recycling microdomains.^9,10,16,17^ This suggested that recycling microdomain components might directly cross-regulate components of the degradative microdomain to maintain the sorting functions of the endosome.

We focused recent studies on understanding the relationship between recycling and degradative microdomains, studying their interplay most extensively in the *C. elegans* coelomocytes. These scavenger-like cells contain unusually large sorting endosomes, often 1 to 5 μm in diameter, allowing direct visualization of endosomal microdomains *in vivo* in the context of the live animal. In these cells, RME-8/SNX-1-marked recycling microdomains clearly lie adjacent to an ESCRT-0 (HGRS-1/HRS marked) degradative microdomain during the early-to-late endosome transition.^8,18^

In *rme-8* and *snx-1* mutants, HGRS-1/HRS over-accumulates on the endosomal membrane at the expense of the unassembled cytoplasmic pool, spreading to fully cover the endosomal membrane and inappropriately encroaching into the recycling microdomains where it is normally absent.^8^ Current models posit that RME-8 in the recycling microdomain recognizes degradative microdomain machinery such as HGRS-1/HRS and activates the Hsc70 folding chaperone to release invading components into the cytoplasm from which they can reassemble elsewhere. Loss of *rme-8* disrupts this balance, allowing degradative machinery to over-assemble on the endosomal membrane, spreading into the recycling microdomain and disrupting cargo sorting. This perturbation leads to commonly recycled proteins such as MIG-14/Wls failing to recycle and increasing trafficking to the lysosome.^8,13,14^ These studies established that endosomal microdomain separation is an actively maintained feature of sorting endosomes. Importantly, the defect in microdomain separation observed upon loss of RME-8 or SNX-1 is not complete, suggesting that additional factors must act to preserve this organization.

To identify additional proteins that maintain microdomain separation and cargo sorting, we performed RME-8 proximity-based biotinylation screens in both *C. elegans* and human cells. These screens identified established recycling microdomain proteins as well as new components that clarify the mechanisms used by the cell to separate recycling and degradative functions. In this study, we focused our efforts on the conserved integral membrane protein SCM-1, a homolog of human SCAMP1 and SCAMP3 (Figures 1A, 1B).^19^ SCAMPs are particularly compelling as potential microdomain regulators due to their extensive self-assembly and interactions with other trafficking machinery, and because cell culture studies have implicated them in both recycling and degradation.^20–24^ However, their precise function remains a subject of debate: while some evidence suggests SCAMPs promote cargo recycling by competing for ESCRT machinery,^25^ subsequent reports suggest they facilitate degradative sorting by promoting intraluminal vesicle formation.^26^

**Fig. 1.**
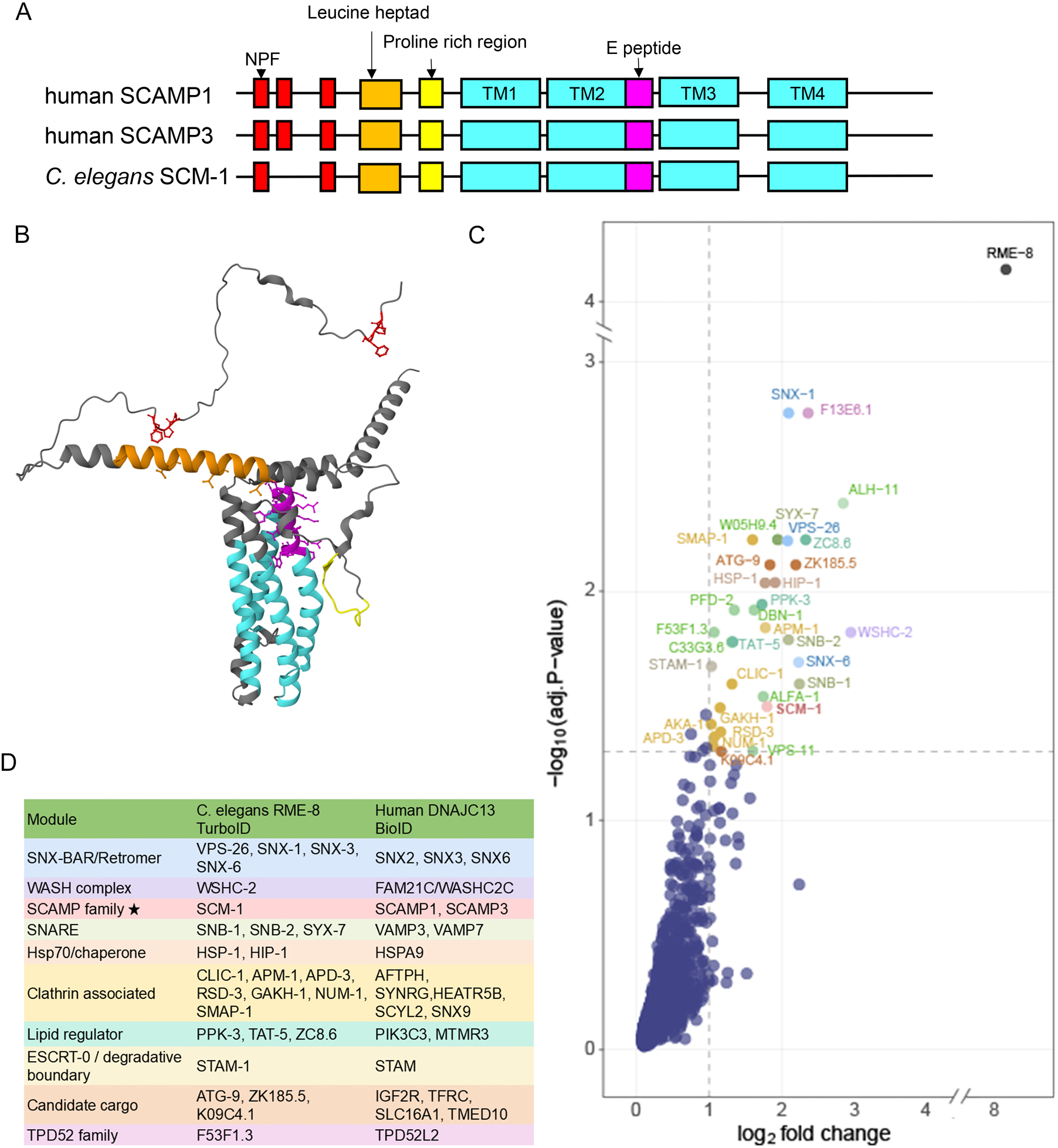
RME-8 proximity-dependent biotinylation identifies SCM-1/SCAMP in the recycling microdomain (A) Domain organization of human SCAMP1, SCAMP3, and C. *elegans* SCM-1. SCM-1 contains two N-terminal NPF motifs (red), whereas human homologs contain three. All SCAMP proteins contain four transmembrane domains (TM, teal). (B) AlphaFold-predicted structure of SCM-1 highlighting NPF motifs (ref), leucine heptad (orange), praline-rich region (yellow), and transmembrane domains (teal). (C) Volcano plot showing RME-8::TurbolD proximity-dependent biotinylation in C. *elegans* intestinal cells versus unfused TurbolD specificity control. x-axis, 109_2_ enrichment over control; y-axis, - log_10_(limma-adjusted P-value). Dashed lines indicate cutoffs (109_2_ fold change> 1, adjusted P < 0.05; 36 significant hits). Significant proximity neighbors are color-coded by functional module as summarized in panel D. SCM-1 is shown in red. Known direct interactors SNX-1 and HSP-1 are among the high-confidence hits.(D) Comparison chart of common endocytic modules that were identified in both proximity labelling experiments.

To understand potential contributions of SCAMP proteins to endosomal microdomains and reconcile conflicting observations regarding their function in endosomal sorting, we investigated the role of SCM-1, the only *C. elegans* SCAMP, within the *C. elegans* coelomocyte system. We find that SCM-1 is largely restricted to endosomal recycling microdomains and is required to maintain their spatial separation from degradative domains through a mechanism distinct from that of RME-8. Consistent with a role in preserving microdomain integrity, *scm-1* mutants inappropriately sort retromer cargo MIG-14/Wls and recycling v-SNARE SNB-2/VAMP3/cellubrevin into the lysosomal pathway, with SNB-2 itself also required for MIG-14 recycling. Importantly, we also identified impaired degradative activity toward transmembrane cargo in *scm-1* mutants, indicating dual requirements for SCM-1 in recycling and degradation. Together, our results define SCM-1 as a key regulator of microdomain boundaries, ensuring the fidelity of both recycling and degradative trafficking pathways.

## Results

### RME-8 proximity-dependent biotinylation identifies SCM-1/SCAMP in the recycling microdomain

We previously identified the J-domain protein RME-8 as a key regulator of endosomal microdomain separation.^8^ RME-8 maintains domain identity by actively removing degradative machinery from the recycling microdomain. Thus, loss of RME-8 leads to degradative component over-accumulation and the intermixing of normally distinct microdomains. However, this mixing is incomplete, suggesting that additional mechanisms must also contribute to maintaining the spatial organization of these functionally opposing domains.

To better understand the composition of the recycling microdomain and identify additional proteins required for microdomain separation *in vivo*, we performed proximity biotinylation experiments in two systems; the *C. elegans* intestine using an RME-8::TurboID fusion, and HEK293 Flp-In T-REx cells using a fusion of BirA* to hRME-8/DNAJC13. Controls included four biological replicates of unfused intestine-specific TurboID in *C. elegans,* and BirA*-FLAG-only and triple-FLAG-only expressing cell lines in HEK293, with the eight HEK293 control samples combined for SAINT analysis.^27^ Proximity-dependent biotinylation is an approach that can identify close neighbors of proteins of interest *in vivo*, biotinylating free primary amines within approximately 10 nm of the fusion protein.^28^ We used streptavidin beads to isolate biotinylated proteins from lysed *C. elegans* and human cells under standard conditions and analyzed captured proteins by mass spectrometry.

The robustness of our proximity labeling datasets is supported by our recovery of well-characterized RME-8/DNAJC13 physical interactors in both species (Figure 1C).

Complete lists of significant proximity neighbors from both screens are provided in Tables S2 (*C. elegans*, limma adjusted P < 0.05, 36 hits) and S3 (HEK293, SAINT score ≥ 0.80, 65 hits), with all primary statistics (fold-change, log-odds, spectral counts where applicable). Among the conserved hits, we recovered *C. elegans* sorting nexin SNX-1 and its human homolog SNX2, along with their heterodimeric partners SNX-6 and SNX6, and the orthologous WASH complex subunit WSHC-2 and FAM21C, proteins with known direct physical interactions with RME-8 or hRME-8/DNJC13 that validate the spatial accuracy of our approach.^13,29,30^ The *C. elegans* screen also recovered retromer component VPS-26, and the HEK293 screen recovered SNX3, consistent with the established role of RME-8 in retrograde recycling.^13,14^

We and others previously established that, as a J-domain co-chaperone, RME-8 recruits Hsc70 to endosomal membranes where it facilitates endosomal coat dynamics.^31–33^ Importantly, our proximity proteome data clearly identified Hsc70 homologs *C. elegans* HSP-1 and human HSPA9. Both screens recovered clathrin associated proteins along with ESCRT-0 subunit STAM-1/STAM, consistent with the known role of RME-8 in disassembly of endosomal clathrin and ESCRT-0.^8,13,14^ Notably, no other ESCRT-associated machinery was recovered.

Among the conserved hits appearing in both proximity proteomes, we were struck by members of the SCAMP family, *C. elegans* SCM-1 and its human homologs SCAMP1 and SCAMP3, integral membrane proteins with no prior connection to RME-8 function (Figure 1A–C). We also recovered homologous SNARE proteins *C. elegans* SNB-2/SNB-1 and human VAMP3 as additional conserved integral membrane components of the RME-8-proximal recycling microdomain. Given the known association of SCAMPs with post-Golgi and endosomal trafficking^19,34^, we prioritized SCM-1 for functional analysis. We also examined the functional relationship between SCM-1 and the co-recovered SNAREs below.

SCAMP proteins have several properties suggesting that they could act as organizers of membrane microenvironments. SCAMPs are known to oligomerize, have been reported to associate with SNAREs, and have been proposed to modulate local lipid composition.^22–24,35–37^ In the context of endosomal sorting, these properties are well suited to potentially support the formation or stabilization of recycling microdomains, where precise coordination of membrane curvature, protein crowding, and cargo selection is likely required. Since SCM-1/SCAMP and the SNAREs identified are broadly expressed in *C. elegans*, we focused most of our analysis on coelomocyte cells, utilizing the clear advantages they provide for microdomain analysis *in vivo*.

### SCM-1 localizes to Golgi and endosomal recycling microdomains

To define the subcellular localization of the only *C. elegans* SCAMP family protein SCM-1, we compared the localization of SCM-1 with markers for several membrane compartments within coelomocytes (Figures 2A-2D”). Early endosomes in coelomocytes typically reside at the cell periphery and move inward as they mature into late endosomes and lysosomes (Figure 2E). SCM-1 localized to small cytoplasmic puncta and partially labeled large ring-like structures. The SCM-1-labeling on ring structures colocalized well with RME-8, identifying the SCM-1 marked structures as endosomal recycling microdomains, consistent with our proximity biotinylation results (Figures 2A-2A“, quantified in 2F). SCM-1 showed minimal colocalization with the HGRS-1/HRS on the same endosomes, indicating a high degree of exclusion from degradative microdomains (Figures 2B-2B”, quantified in 2F). The small puncta co-labeled with Golgi marker AMAN-2 identifying them as dispersed Golgi ministacks as are typical of invertebrate cells (Figures 2C-2C“, quantified in 2F).^38^ Note that adjacency of the SCM-1 and AMAN-2 suggests that SCM-1 may be enriched at the trans-Golgi, since AMAN-2 is cis-Golgi enriched (Figure 2E).^39^ SCM-1-labeling was also absent from LMP-1-marked lysosomes (Figures 2D-2D”). We conclude that SCM-1 localizes predominantly to Golgi ministacks and RME-8-positive endosomal recycling microdomains, where it is largely excluded from degradative ESCRT-0 domains, indicating selective enrichment within the recycling membrane microdomain.

**Fig. 2.**
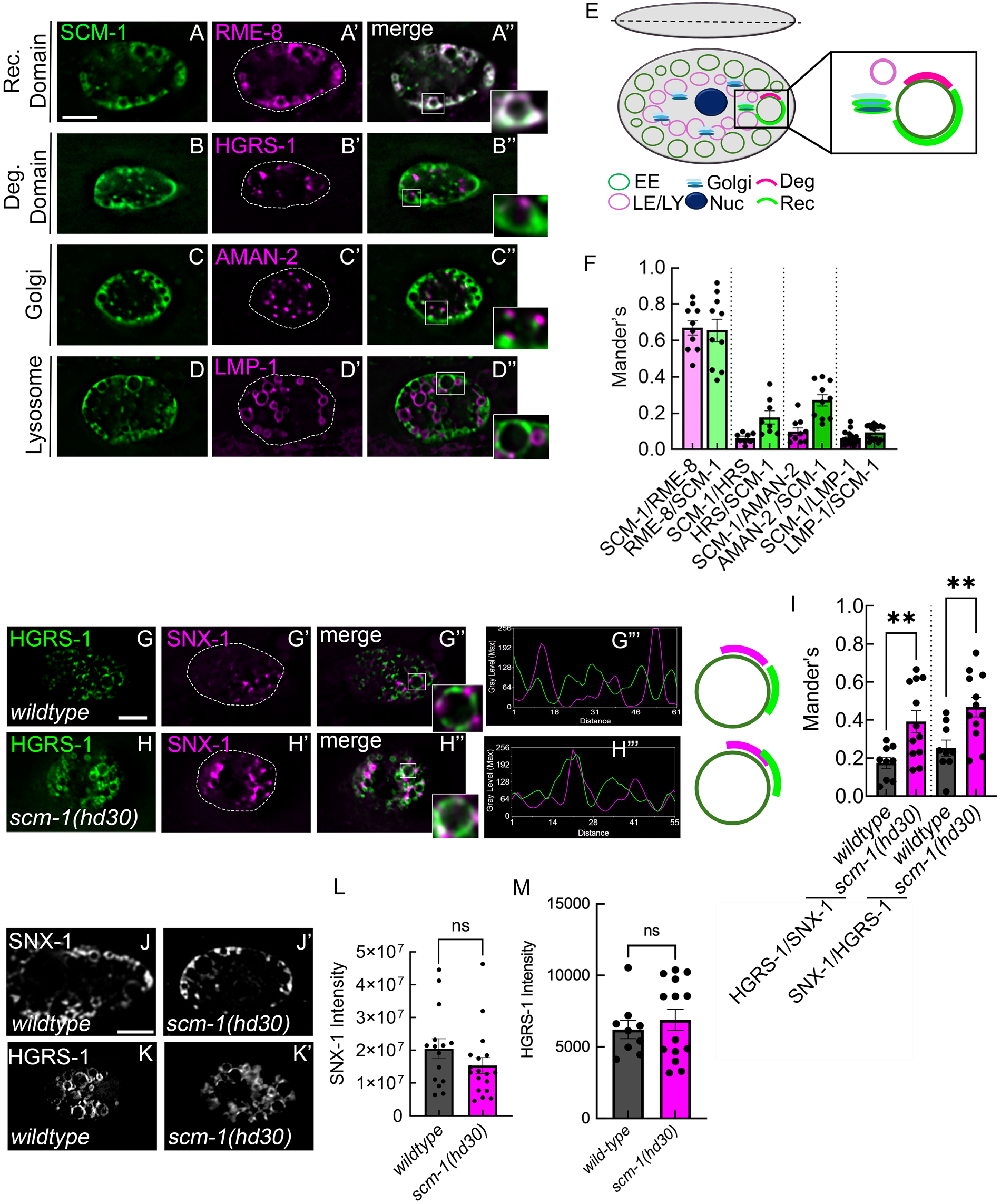
SCM-1 localizes to recycling microdomains and is required to maintain microdomain separation (A-D) Confocal images of GFP::SCM-1 in coelomocytes with markers for endosomal recycling microdomains (tagRFP::RME-8), degradative microdomains (tagRFP::HGRS-1), Golgi (AMAN-2::tagRFP), and lysosomes (LMP-1::GFP) with mScarlet::SCM-1. SCM-1 localizes to endosomal membranes where it shows strong overlap with RME-8, minimal overlap with HGRS-1, and is largely excluded from lysosomes, while also labeling Golgi ministacks. Insets are 2-fold larger. (E) Schematic showing Z-slice of a coelomocyte with localization of organelles. (F) Quantification of colocalization using Manders’ coefficients. (G-H) Recycling (tagRFP::SNX-1) and degradative (citrine::HGRS-1) microdomains remain spatially distinct in wild type but exhibit increased overlap in scm-1(hd30) mutants. (G“-H”) Line scan analysis along the endosomal limiting membrane reveals loss of sharp domain boundaries in mutants. (I) Quantification of increased overlap between SNX-1 and HGRS-1. (J-K) Total endosomal levels of SNX-1 and HGRS-1 are unchanged in scm-1 mutants. (L-M) Quantification of total intensity levels of SNX-1 and HGRS-1. Data are presented as mean ± SEM. Each data point representing a single coelomocyte and data shown (n>12) is representative of three independent experiments. Scale bar: 5 µm (A-D, G-H, J-K). ** p:s; 0.01 by Student’s t-test (I)

### SCM-1 is required to maintain normal separation of endosomal recycling and degradative microdomains

Given the established role of RME-8 in regulating endosomal microdomain organization, we investigated whether SCM-1 similarly contributes to this spatial architecture. To address this, we utilized the previously characterized deletion allele *scm-1(hd30)*^40^, which removes the entire fourth exon and parts of the flanking introns. Because exon four encodes all four predicted transmembrane domains, *hd30* is expected to be a functional null.

To test SCM-1 for influence on microdomain structure we measured the overlap of recycling microdomain marker SNX-1 and degradative microdomain marker HGRS-1/HRS, comparing control and *scm-1(hd30)* mutant animals (Figures 2G-2H). Line-scan analysis of fluorescence intensity along the endosomal circumference revealed that recycling and degradative microdomains still assemble in *scm-1* mutants but fail to remain distinct (Figure 2G’’’ and H’’’). We quantified this loss of separation using Manders’ coefficients and found a significant increase in the overlap between HGRS-1 and SNX-1 signals in the absence of SCM-1 (Figure 2I). Collectively, these results demonstrate that while SCM-1 is dispensable for the formation of endosomal microdomains, it is essential for maintaining their spatial segregation.

### SCM-1 affects microdomains differently than does RME-8

We next asked whether SCM-1 enforces microdomain separation through similar mechanisms to those we previously established for RME-8. In *rme-8* mutants, ESCRT-0 component HGRS-1/HRS spreads around the endosomal membrane, accumulating on the endosomal membranes from the cytoplasm. This results in an increased average intensity of HGRS-1 labeling despite the spread over a larger membrane area. At the same time SNX-1 does not expand around the endosome in *rme-8* mutants but instead displays increased accumulation within recycling microdomains with reduced dynamics.^8,18^

Unlike in *rme-8* mutants, in *scm-1* mutants we did not find an increased intensity of HGRS-1 or SNX-1 (Figures 2J-2K’, quantified in 2L-2M) when compared to controls, indicating that despite abnormal microdomain overlap, the level of recruitment of these proteins to the endosomal membrane is unchanged. Taken together, these data show that while SCM-1 and RME-8 are both important for the separation of discrete microdomains, the mechanisms by which they contribute are distinct.

### *scm-1* mutants missort recycling cargo MIG-14

Given that SCM-1 resides in the recycling microdomain and is required for the correct partitioning of microdomains, we examined whether recycling cargo proteins were properly sorted in the absence of SCM-1. We focused on the well-characterized retrograde recycling cargo MIG-14/Wls. At the plasma membrane, MIG-14 is internalized via clathrin-mediated endocytosis into early/sorting endosomes. From there, it is recycled to the Golgi through a retromer and RME-8-dependent pathway, preventing wholesale entry into the degradative pathway. In retromer or *rme-8* mutants, MIG-14 is missorted into late endosomes/lysosomes, where active degradation leads to lower MIG-14 protein levels.^8,13,41^

Interestingly, we found an expanded population of MIG-14 positive vesicles in *scm-1* mutants without any change in MIG-14 average intensity (Figures 3A-3B, quantified in 3D-3E, illustrated in 3C). To determine if this reflected a change in the distribution of MIG-14 among endolysosomal vesicle types, we measured MIG-14 colocalization with endolysosomal markers *in vivo*. In control animals, MIG-14 colocalizes well with RAB-5-labeled early endosomes, but very little signal is found in RAB-7 or LMP-1 labeled late endosomes and lysosomes (Figures 3F-F”, 3H-H”, 3J-J”), consistent with efficient recycling and rescue of MIG-14 from the degradative pathway. In *scm-1* mutant animals, however, while a population of MIG-14 still colocalizes with early endosomes, its localization to the limiting membrane of both late endosomes and lysosomes is significantly increased (Figures 3G-G”, 3I-I”, 3K-K”, illustrated in 3L, quantified in 3M). In summary, while MIG-14 is normally efficiently recycled away from the degradative pathway, loss of SCM-1 causes misrouting of MIG-14 to late endosomes and lysosomes, indicating a recycling defect. The retention of MIG-14 on the lysosomal limiting membrane and maintained overall MIG-14 level could indicate an additional defect in ESCRT-mediated degradation, as previously observed upon perturbation of microdomain boundaries in *ubc-13* mutants, which display a dual recycling and degradative defect.^42^

**Fig. 3.**
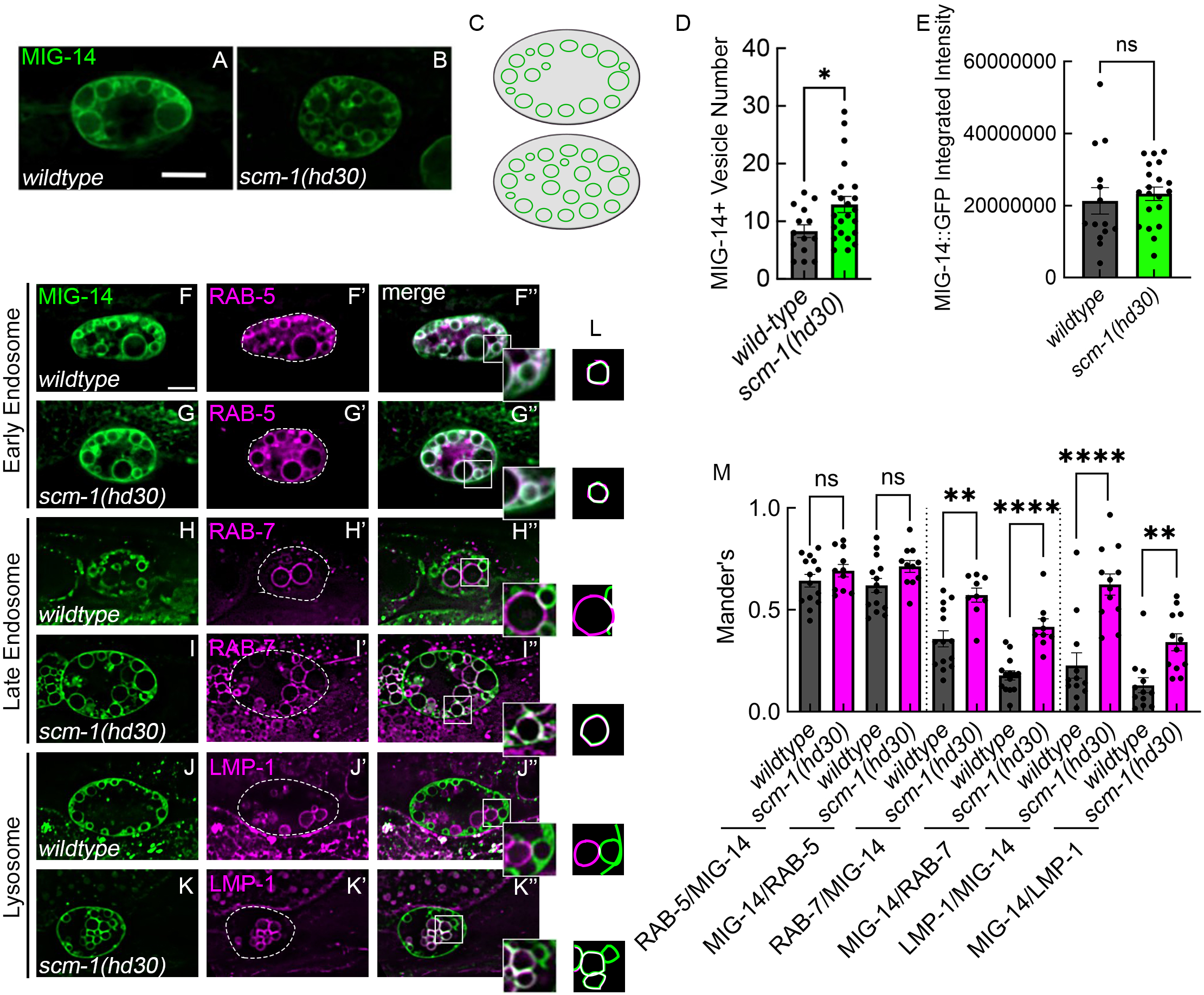
*scm-1* mutants misroute recycling cargo MIG-14 to late endosomes and lysosomes. (A-B) MIG-14::GFP localization in wildtype and *scm-1(hd30)* coelomocytes. MIG-14 accumulates in internal vesicles in mutants. (C) Schematic of altered cargo distribution. (D-E) Quantification of vesicle number and total fluorescence intensity. (F-K) Colocalization of MIG-14::GFP with endomembrane markers tagRFP::RAB-5, mScarlet::RAB-7 and LMP-1::tagRFP. Insets are 2-fold larger. (L) Cartoons of insets from F-K (M) Quantification by Manders’ correlation coefficient shows increased localization of MIG-14::GFP to mScarlet::RAB-7 and LMP-1::tagRFP positive late endosomal and lysosomal compartments. Data are presented as mean± SEM. Each data point representing a single coelomocyte and data shown (n>9) is representative of three independent experiments. Scale bar: 5 µm (A-B, F-K). *p::; 0.05, ** p::; 0.01, ****p::; 0.0001 by Student’s t-test (D, M).

### SNB-2 colocalizes with SCM-1 on early endosomes

Having established that the loss of SCM-1 disrupts the recycling of the canonical retromer cargo MIG-14, we next sought to identify other RME-8-proximal factors that might contribute to SCM-1-associated recycling. Among the conserved proteins identified in our proximity datasets, the SNAREs SNB-1 and SNB-2 and their human homolog VAMP3/cellubrevin were particularly notable, given that VAMP3-family proteins are well-characterized for their roles in endocytic recycling.^6,43–45^ Our datasets also indicated enrichment of SNAREs *C. elegans* SYX-7 and human VAMP8. Together, these observations suggested that a specific subset of SNARE proteins might function with SCM-1 to facilitate the efficient sorting of transmembrane cargo.

To test whether any of these SNAREs could act with SCM-1 *in vivo*, we asked whether GFP-tagged SNB-2, SNB-1, SYX-7, or VAMP-7 (homolog of human VAMP8) were enriched in the SCM-1-positive recycling microdomain in coelomocytes (Figures 4A-4D”, illustrated in 4E, quantified in 4F). SNB-2, SNB-1, and SYX-7 showed a high degree of colocalization with SCM-1 on endosomes. VAMP-7 showed low colocalization with SCM-1 (Figures 4D-4D“, illustrated in 4E, quantified in 4F). Together, these experiments identify SNB-2, SNB-1, and SYX-7, as recycling-microdomain-associated SNAREs that are positioned as potential contributors to SCM-1-dependent trafficking.

**Fig. 4.**
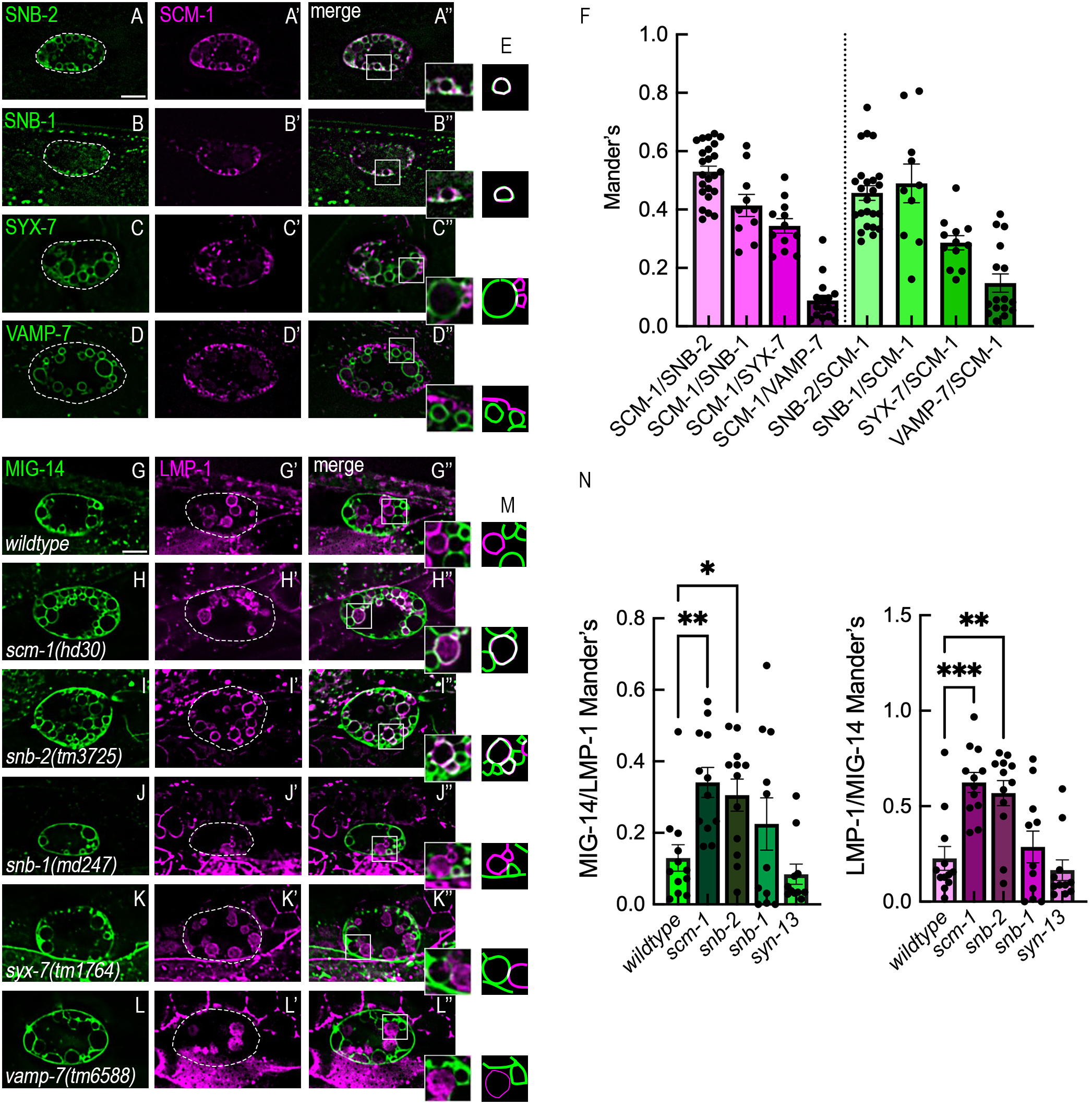
SNB-2 localizes to recycling microdomains and phenocopies *scm-1* cargo-sorting defects. (A-D) Colocalization of mScarlet::SCM-1 with GFP-tagged SNAREs. SNB-2 and SNB-1 show the highest degree of overlap with SCM-1, with partial overlap observed for SYX-7, and minimal overlap for VAMP-7. Insets are 2-fold larger. (E) Cartoons of insets from A-D. (F) Quantification of colocalization by Manders’ correlation coefficient. Each data point represents a single coelomocyte with n>9 animals. (G-L) MIG-14 localization in wild type and SNARE mutants. (M) Cartoons of insets from G-L. (N) snb-2 mutants show increased colocalization of MIG-14 with LMP-1, phenocopying the scm-1 sorting defect, whereas other SNARE mutants do not. Data are presented as mean± SEM. Each data point represents the average of one independent experiment with coelomocytes n>12. Scale bar: 5 µm (A-D, G-L). no annotation indicated no signifigance, *p::5 0.05, **p::5 0.01, ***p::5 0.001 (one-way ANOVA) (N). Scale bar is 5microns.

### *snb-2* phenocopies key aspects of the *scm-1* cargo-sorting and endolysosomal phenotypes

We predicted that loss of any SNAREs that contribute to SCM-1-dependent recycling should phenocopy at least part of the *scm-1* cargo-sorting defect. To test this, we assayed for missorting of MIG-14 into the LMP-1 marked lysosomal compartment, comparing controls with *snb-2(tm3725), snb-1(md247), syx-7(tm1764),* and *vamp-7(tm6588)* loss-of-function mutants (Figures 4G-4L”, illustrated in 4M, quantified in 4N). Importantly, we observed a substantial increase in colocalization between MIG-14 and LMP-1 in the *snb-2* mutant, strongly phenocopying the *scm-1* mutant sorting defect (compare Figures 4H” to 4I”). None of the other SNARE mutants affected MIG-14 sorting, emphasizing the specificity of the SNB-2 effect.

We also noted additional effects on the endolysosomal system that further parallel the *scm-1* phenotype. In *snb-2* mutants, lysosomes were more numerous, consistent with the increased lysosome number observed in *scm-1* animals (Figures S1A, quantified in S1B, S1D). This result suggests recycling is reduced, leading to enhanced delivery of cargo to the degradative pathway. *scm-1* mutants also have smaller lysosomes (Figure S1C). In contrast, *snb-1* mutants displayed an overall reduction in the size of both MIG-14-positive endosomes and lysosomes, a phenotype similar to that previously observed in the coelomocytes of mutants with decreased endocytosis from the plasma membrane (Figure S1E).^46,47^ Together, these data emphasize that *snb-2* and *scm-1* mutants share common phenotypes, likely reflecting reduced recycling and consequent rerouting of cargo to lysosomes. *snb-1* mutants, however, exhibit a global reduction in the number and size of endolysosomal compartments, indicating a mechanistically distinct defect.

### SCM-1 regulates SNB-2 recycling

The phenocopy of the MIG-14 sorting defect in *snb-2* mutants suggested that SNB-2 functions with, or downstream of, SCM-1 in recycling microdomain activity. If SCM-1 controls SNB-2 sorting, we would expect SNB-2 localization to be altered in an *scm-1* mutant. Indeed, we noted that in *scm-1* mutants, SNB-2 labeling was reduced on peripheral endosomes that are LMP-1-negative, while displaying increased overlap with LMP-1 in lysosomes (Figures 5A-5B”, illustrated in 5C). Consistent with this, Manders’ coefficients revealed a significant increase in SNB-2 and LMP-1 colocalization in the *scm-1* mutant background, indicating that SNB-2 is missorted to lysosomes in the absence of SCM-1 (Figure 5D). Loss of SNB-2 from early endosomes is likely to contribute to failed recycling in *scm-1* mutants.

**Fig. 5.**
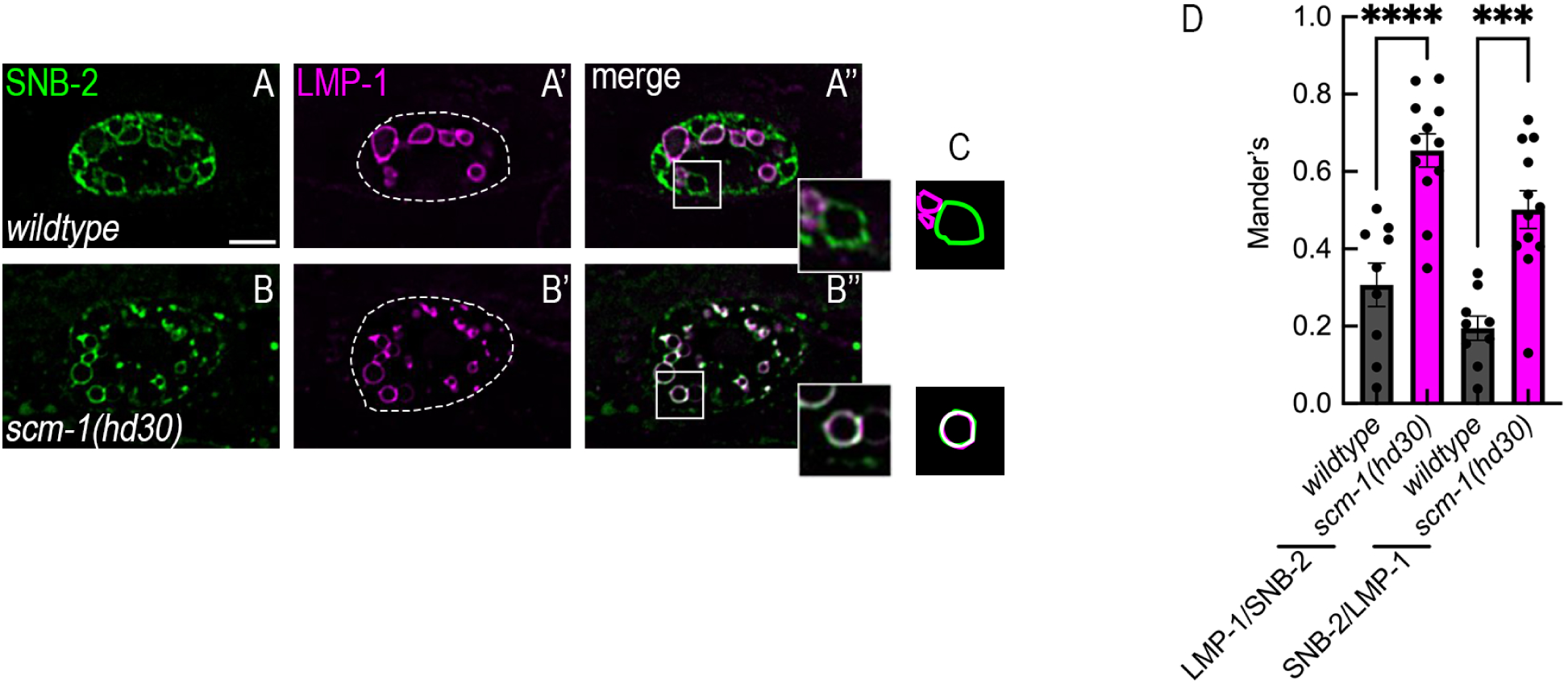
SCM-1 regulates SNB-2 sorting. (A-8) Localization of SNB-2 in wild-type and *scm-1(hd30)* coelomocytes. GFP::SNB-2 is reduced on peripheral endosomes and accumulates in LMP-1::mKate2-positive lysosomes in scm-1 mutants. Insets are 2-fold larger. (C) Cartoons of insets from A-8. (D) Quantification by Manders’ correlation coefficient shows increased localization of SNB-2 to lysosomes in *scm-1* mutants. Data are presented as mean± SEM. Each data point representing a single coelomocyte and data shown (n>12) is representative of three independent experiments. Scale bar: 5 µm (A-8). ***p::; 0.001, ****p::; 0.0001 by Student’s t-test (D)

### SCM-1 affects degradation of transmembrane cargo

While our MIG-14 and SNB-2 data strongly support a role for SCM-1 in recycling microdomain function, our results show relatively normal overall levels of MIG-14, and accumulation of MIG-14 and SNB-2 on the lysosome limiting membrane in *scm-1* mutants. Along with previous work on mammalian SCAMP proteins, these results suggested a potential role of SCM-1 in membrane protein degradation. Loss of microdomain separation might be expected to perturb not only recycling, but also degradative sorting of membrane cargo. We therefore tested whether SCM-1 is required for efficient transmembrane cargo degradation in an independent assay, leveraging a well-characterized trafficking transition in the *C. elegans* germ line. During ovulation and passage through the spermatheca, cortical granules undergo exocytosis and contribute membrane proteins to the oocyte surface. Following fertilization, a subset of these surface proteins is rapidly ubiquitylated, internalized, and delivered to endosomes for ESCRT-mediated delivery to intraluminal vesicles and subsequent lysosomal degradation.^48,49^ Normally, CAV-1::GFP is efficiently degraded such that embryos at the two-cell stage and beyond contain little detectable fluorescence.^49,50^ We found that in *scm-1(hd30)* mutants, embryos retained significantly higher CAV-1::GFP signal compared to controls (Figures 6A-6B’, quantified in 6C), indicating delayed degradation. These data indicate a requirement for SCM-1 in efficient degradation of transmembrane cargo, in addition to its role in recycling.

**Fig. 6.**
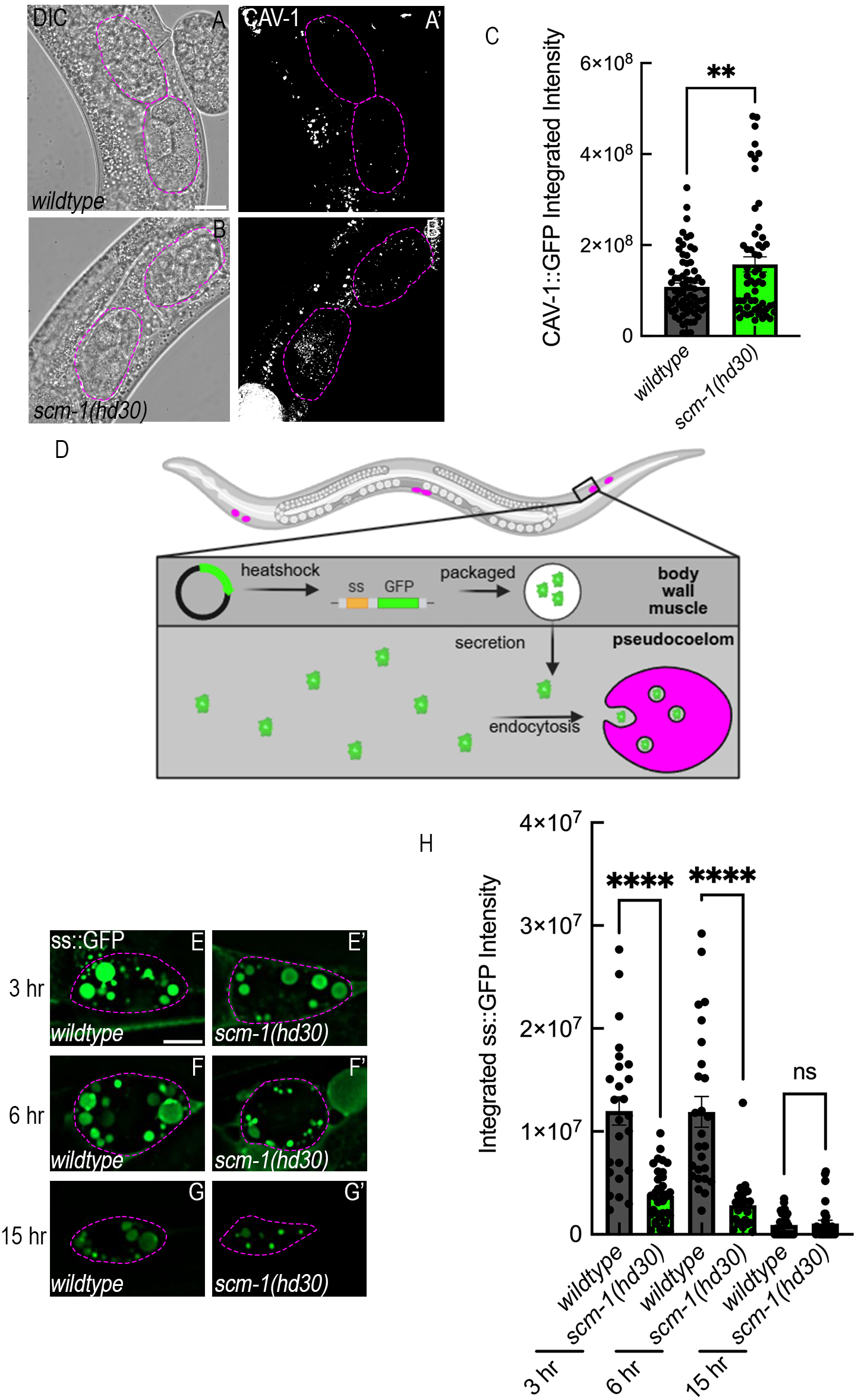
SCM-1 is required for efficient degradation of transmembrane cargo but not for bulk endolysosomal function. (A-B) CAV-1::GFP in embryos from wild-type and *scm-1(hd30)* animals. CAV-1 persists to later stages in scm-1 mutants. Region of interest (embryo, pink dotted outline) was drawn in DIC and transferred to GFP images to measure total GFP intensity of embryos at 2-cell stage and later. (C) Quantification of CAV-1::GFP intensity indicates delayed degradation. Data are presented as mean± SEM. Each data point represents a single embryo with total animals n>12. (D) Schematic of heat-shock-driven secreted GFP (ssGFP) uptake assay. (E-G) Coelomocyte uptake of ssGFP over time. Wildtype and *scm-1(hd30)* worms were heat-shocked at 34°C for 1 hr and rested at 20°C for 3,6-, and 15-hours. **(H)** Quantification shows similar kinetics of GFP clearance despite reduced uptake in mutants. All worms were grown at 25 °C. Each data point representing a single coelomocyte and data shown (n>12) is representative of three independent experiments **(H).** Scale bar: 5 µm (A, B, E-G) ** p:5 0.01, ****p:5 0.0001 by Student’s t-test (C, **H).**

### SCM-1 does not affect endosomal maturation, acidification, or fluid phase transport from endosome to lysosome

The retention of MIG-14 on lysosomal membranes and the delay in CAV-1 degradation are consistent with an SCM-1 effect on the efficiency of ESCRT-mediated degradation, but these phenotypes could also reflect a general defect in endolysosomal maturation or degradative capacity. To help distinguish between these possibilities, we systematically assayed for evidence of defective endosomal maturation, lysosome acidification, and bulk delivery to lysosomes. Defects in maturation can stall cargo in intermediate endosomes positive for both RAB-5 and RAB-7.^51,52^ However, colocalization of GFP::RAB-5 and endogenously tagged mScarlet::RAB-7 was unchanged in *scm-1* mutants (Figure S2A, illustrated in S2B, quantified in S2C). Similarly, colocalization of RAB-7 and LMP-1 remained unchanged (Figure S2A, illustrated in S2B, quantified in S2C), indicating that SCM-1 does not broadly impair endosomal maturation or late endosome-lysosome fusion.

We evaluated lysosome acidification using a *C. elegans*-optimized ratiometric FIRE-pHLy pH biosensor.^53^ Despite the morphological changes observed in *scm-1* lysosomes, we measured equivalent mTFP1/mCherry fluorescence ratios between controls and mutants, indicating that *scm-1* mutant lysosomes are properly acidified (Figure S2D, quantified in S2E).

To evaluate bulk endocytic delivery to lysosomes, we utilized two assays: Cy5-BSA microinjection into the body cavity (pseudocoelom), and heat shock driven expression of signal-sequence GFP (ssGFP) secreted into the body cavity. We observed that BSA uptake and delivery kinetics to the LMP-1 marked lysosomal compartment of coelomocytes were equivalent at both 5-and 30-minute timepoints, with almost all lysosomes becoming BSA positive by 30 minutes in both control and mutant animals (Figure S2F, quantified in S2G). In the complementary heat shock driven ssGFP assay, we noted a reduction in ssGFP uptake similar to that observed in *rme-8* mutants (Figures 6E-6G, illustrated in 6D, quantified in 6H).^31^ We attribute the reduced GFP uptake to probable recycling defects in a scavenger receptor known to affect GFP but not BSA uptake (Hanna Fares, personal communication).^54^ GFP signal within coelomocytes at the 15-hr timepoint was similar to control, consistent with a relatively normal lysosomal degradative capacity. Collectively, these data indicate that endosomal maturation and lysosomal degradative capacity remain largely unaffected by SCM-1, supporting a specific role for SCM-1 in the sorting of transmembrane cargo at the sorting endosome.

## Discussion

Endosomal sorting depends on the spatial segregation of competing trafficking machineries into physically distinct microdomains on the endosomal limiting membrane. While the molecular composition of key sorting complexes, including Retromer and ESCRT, has been extensively defined^2,5–7^, how these complexes are organized into discrete membrane domains, and how the composition and identity of those domains are established and maintained, remain much less well understood. Prior work demonstrated that recycling and degradative activities occur in adjacent but distinct regions of the same endosome^8,18^, yet the mechanisms preserving this spatial segregation remain unclear. In this study, we identify the conserved tetraspan protein SCM-1/SCAMP as a key regulator of endosomal microdomain boundary integrity and show that it functions through a mechanism distinct from the previously defined RME-8 pathway.

### Multiple mechanisms enforce microdomain separation

Our previous work established that the J-domain protein RME-8 promotes microdomain separation by regulating the assembly/disassembly dynamics of degradative machinery. In *rme-8* mutants, ESCRT-0 components such as HGRS-1/HRS accumulate on the endosomal membrane and spread into recycling microdomains^8^, consistent with a failure to remove inappropriately assembled degradative complexes. This was a unique property of RME-8 and its partners SNX-1/SNX-6 and did not occur in mutants lacking other retromer-associated proteins such as VPS-35 or SNX-3.^8^ These findings support a model in which RME-8, acting with Hsc70, promotes the disassembly or release of invading components, thereby preserving domain identity.

In contrast, loss of *scm-1* produces a distinct phenotype. Recycling and degradative markers still assemble into recognizable domains, but these domains fail to remain spatially distinct and instead display irregular intermixing. We do not observe increased membrane association of either HGRS-1 or SNX-1 in *scm-1* mutants, indicating that SCM-1 does not primarily regulate the abundance or turnover of these complexes on the membrane. Rather, SCM-1 appears to function at the level of domain organization, maintaining the spatial separation of pre-assembled sorting machineries.

Taken together, these observations support a model in which endosomal microdomain integrity is maintained by at least two complementary processes: (1) active removal of inappropriate components, mediated by RME-8 and associated chaperone activity, and (2) intrinsic stabilization of domain identity, mediated by membrane-resident factors such as SCM-1. Importantly, these findings support the view that adjacent microdomains do not function independently but instead engage in reciprocal cross-regulation to balance opposing sorting activities.^12^ Such spatially constrained cross-regulation provides a flexible means to maintain sorting fidelity while allowing rapid adjustment of trafficking outputs in response to cellular conditions.

### SCM-1 maintains domain identity rather than directly regulating sorting complexes

Previous studies of mammalian SCAMP proteins have yielded conflicting conclusions, with some reports implicating SCAMPs in recycling and others in degradative sorting.^25,26^ Our data argue against a primary role for SCM-1 in directly regulating ESCRT function. SCM-1 is strongly enriched in recycling microdomains and largely excluded from degradative regions, and lacks canonical ESCRT-interacting motifs, such as PSAP sequences that mediate engagement with ESCRT-I components.

Instead, our findings support an indirect but central role for SCM-1 in maintaining the identity of the recycling domain. In *scm-1* mutants, loss of spatial separation is accompanied by defects in both recycling and degradation of transmembrane cargo. The retromer cargo MIG-14/Wls is misrouted to late endosomes and lysosomes^13,41^, yet this cargo does not appear to be efficiently degraded, and an independent degradative cargo, CAV-1, shows delayed turnover.^49,50^ These defects occur in the absence of detectable changes in endosomal maturation, lysosomal acidification, or bulk fluid-phase trafficking (Figure 7).

**Fig. 7.**
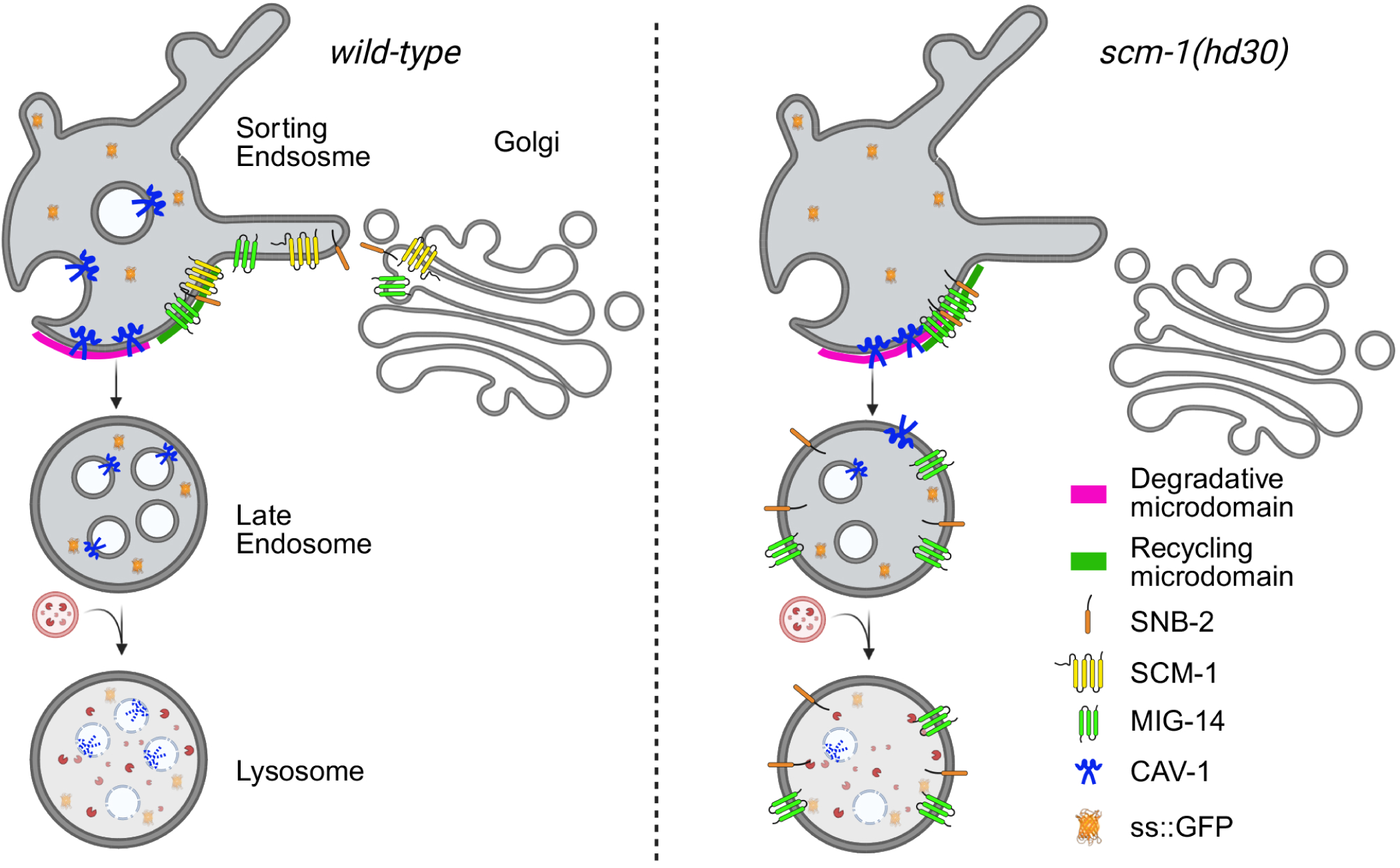
Model for SCM-1 function in retrograde cargo sorting and microdomain organization. SCM-1 localizes to recycling microdomains where it maintains spatial separation from degradative ESCRT domains. Loss of SCM-1 results in intermixing of microdomains and misrouting of cargo such as MIG-14 and SNB-2, into the degradative pathway, reducing the efficiency of both recycling and degradation.

Disruption of these boundaries likely permits inappropriate mixing of sorting machineries, reducing the efficiency of both cargo retrieval and degradative sorting. More fundamentally, such mixing may allow direct interference between opposing trafficking systems, a condition that has been proposed to compromise sorting fidelity. In this view, previously reported roles for SCAMP proteins in either recycling or degradation may reflect indirect consequences of altered domain organization rather than direct participation in one pathway or the other.

### A functional relationship between SCM-1 and SNB-2

Our proximity labeling and localization studies identify the v-SNARE SNB-2 as a conserved component of the recycling microdomain. SNB-2 colocalizes with SCM-1, and loss of *snb-2* phenocopies key aspects of the *scm-1* cargo-sorting defect. In *scm-1* mutants, SNB-2 is mislocalized to the lysosome limiting membrane, indicating that its proper distribution depends on SCM-1. These findings suggest that SCM-1-dependent domain organization is required to maintain SNB-2 within the recycling microdomain, which in turn is essential for efficient recycling.

SNARE proteins are best known for their roles in membrane fusion, including the SNB-2 ortholog VAMP3 in endosome to Golgi transport^43^, yet their spatial restriction may also contribute to maintaining compartment identity. Mislocalization of SNB-2 in *scm-1* mutants may therefore reflect a broader failure to preserve the molecular composition of the recycling domain. Consistent with this idea, several SNARE proteins have been proposed to associate with SCAMP family members, although evidence for direct physical interaction remains scant.^23,24,55,56^

### Proximity biotinylation defines recycling microdomain components

Defining the molecular composition of endosomal microdomains *in vivo* has been challenging, as these domains are dynamic, small enough in most cells to confound simple light-based microscopy methods, and not readily separable by conventional biochemical approaches. Proximity-dependent biotinylation provides an effective strategy to overcome these limitations by capturing proteins based on spatial proximity rather than stable complex formation. Using RME-8-directed proximity labeling in both *C. elegans* and human cells, we recovered a highly coherent set of recycling microdomain components, including established SNX-BAR/retromer machinery, WASH complex subunits, and Hsc70 chaperones^8,13,57^, validating the spatial specificity of the approach. Importantly, this strategy also identified conserved factors not previously assigned to recycling microdomains, including SCM-1/SCAMP and SNB-2/VAMP3, highlighting its ability to reveal domain-associated proteins beyond stable complexes.

While proximity labeling captures proteins within a defined radius and does not distinguish direct interactions^58^, this feature is particularly well suited to the study of microdomains, where organization is defined by spatial proximity and dynamic interactions. Together, these results demonstrate that proximity biotinylation is a powerful approach for defining the molecular landscape of membrane microdomains *in vivo*.

### Potential mechanisms for SCM-1-dependent microdomain stabilization

While we do not yet understand precisely how SCM-1 promotes microdomain separation, our finding that neither HGRS-1 nor SNX-1 membrane levels are altered in *scm-1* mutants argues against a role in regulating the assembly dynamics of peripheral coat complexes, distinguishing SCM-1 from the RME-8/Hsc70 uncoating pathway.

SCAMPs are abundant transmembrane components of endomembrane compartments capable of self-association and interaction with multiple trafficking factors.^19,23,34,36,59^ This self-association property suggests that SCM-1 may form membrane-resident assemblies that restrict lateral diffusion and stabilize local protein organization. Such higher-order assemblies are a recurring feature of microdomain-forming systems, including tetraspanins, SNX-BAR polymers, and ESCRT clusters.^60–62^

In addition, diffusion barriers have been proposed to contribute to microdomain integrity, including actin networks associated with recycling domains and ESCRT assemblies within degradative regions.^57,63^ SCAMP proteins have also been implicated in the regulation of membrane lipid composition and organization^35,64,65^, raising the possibility that SCM-1 contributes to lipid environments that favor recycling machinery cohesion, tight molecular packing, and/or disfavors ESCRT assembly.

### Broader implications for organelle organization and physiological control

The identification of SCM-1 as a regulator of microdomain boundaries highlights the importance of higher-order membrane organization in endosomal function. The coexistence of opposing sorting machineries on a shared membrane necessitates mechanisms that preserve spatial separation and prevent functional interference.

Microdomain organization may therefore provide a structural framework for balancing recycling and degradation, allowing cells to dynamically tune trafficking outputs in response to changing physiological demands.^12^ For instance, microdomain organization appears sensitive to cargo load and composition^42,66^, rapidly changing properties *in vivo*, with cargo itself contributing to degradative domain cohesion.

More broadly, these findings suggest that regulation of membrane microdomain identity, distinct from regulation of individual protein complexes, is a central feature of organelle organization. Understanding how such domains are established and maintained through reciprocal mechanisms will be essential for defining the logic of intracellular trafficking systems.

## Materials and Methods

All *C*. *elegans* strains were derived originally from the wild-type Bristol strain N2. Worm cultures, genetic crosses, and other *C*. *elegans* husbandry were performed according to standard methods.^67^ All strains used in this study are listed in Table S1.

### Plasmid construction and transgenic strains

We utilized the *snx-1* promoter (*psnx-1)* to drive broad expression of fluorescent markers in coelomocytes and other tissues. Plasmid constructions utilized Grant lab-modified MiniMos enabled vectors pCFJ1662 (Hygromycin resistant; Addgene 51482) and pCFJ910 (G418 resistant; Addgene 44481; gifts of Erik Jorgensen from University of Utah). Genes of interests are first cloned into Gateway entry vector pDONR221 by BP reaction (Invitrogen). *C. elegans* optimized FIRE-phLY was synthesized with *C. elegans* preferred codon bias and *C. elegans* synthetic introns, and *C. elegans* LMP-1 was substituted for mammalian LAMP1. Then pDONR221 plasmids are combined with pCFJ1662/pCFJ910 vectors with fluorescent protein fused to C-/N- termini by Gateway LR clonase II reaction (Invitrogen). Single copy integrations (pwSi) were obtained by MiniMOS technology.^68^ Low copy integrated transgenic lines (pwIs) were obtained by microparticle bombardment method.^69,70^

### Image acquisition of C. elegans

All worms were grown on nematode growth media (NGM) plates at 20°C, synchronized by picking L4s, and shifted to 25°C overnight to image the next day as young adults.

Live worms were mounted on 8% (wt/vol) agarose pads in 25 mM levamisole solution.

Multiwavelength fluorescence images were taken with 100x oil immersion objective using Zeiss Axiovert Z1 microscope with X-Light V2 Spinning Disk Confocal Unit (CrestOptics), 1-line LDI Laser Launch (89 North), Prime 95B Scientific CMOS camera (Photometrics). Software used for obtaining images was Metamorph 7.10. To capture the entire coelomocyte, Z series of 15-25 optical sections of 0.2 µm increments were selected. Images were deconvolved using AutoDeblur X3 software (AutoQuant Imaging).

### Image analysis

Fluorescence intensity, count, and length measurements were obtained from Metamorph 7.10. Images were calibrated for distance and intensity prior to analysis. Vesicle diameters were drawn using the line tool to denote active regions and distances were quantified using region measurements. For average and integrated intensities, the perimeter of coelomocytes was traced and selected as active region and color channel was manually thresholded prior to quantification. Endosome numbers were obtained by manually counting objects. Colocalization analysis of two colors was obtained with ImageJ (Fiji) software with JaCoP (Just Another Co-localization Plugin) plugin. Region of interest is selected from tracing a coelomocyte from one channel and transferred to the other channel prior to manually thresholding both channels for Manders’.

### BSA microinjections and heat shock assays

Bovine Serum Albumin (BSA) Alexa Fluor™ 647 conjugate was from Invitrogen (A34785). 1 mg/mL BSA was injected into the pseudocoelom; worms were recovered on seeded NGM plates and rested at room temperature for indicated amount of time (5 or 30 minutes). Then, plates were moved onto ice and kept on ice until imaging. For heat shock assays, worms were placed at 34 °C for 30 minutes and placed at 20 °C to rest for indicated amount of time (3, 6, 15 hours) prior to imaging.

### Graphing and statistics

All graphs were made by GraphPad Prism 11 software, with bars representing SEM. Significance was determined by unpaired Student’s t-test or one-way ANOVA with multiple comparisons. Data were considered statistically different at *P* < 0.05. *P* < 0.05 is indicated with single asterisks, *P* < 0.01 with double asterisks, *P* < 0.001 with triple asterisks, and *P* < 0.0001 with quadruple asterisks.

### Proximity biotinylation

The biotinylation-labeling assay in *C. elegans* fused a highly active version of BirA (TurboID) to the C-terminus of RME-8 and expressed the fusion construct as a single-copy integrated transgene driven by *vha-6* intestinal-specific promoter in the AX7884 strain. The AX7884 strain has been previously CRISPR/Cas9 genome edited to add a His_10_ tag to all four endogenously biotinylated carboxylases.^71^ This approach allows for depletion of endogenous biotinylated proteins using immobilized metal affinity chromatography prior to mass spectroscopy, enhancing sensitivity during detection of artificially biotinylated proteins. The biotinylation-labeling assay in human cells used BirA* (R118G) fused to the N-terminus of human RME-8/DNAJC13 expressed in HEK293-FlpIn-TREx cells.

### Growing animals for proteomics analysis

For each biological replicate, *C. elegans* were started as ten 6-cm plates with 6 L4 stage hermaphrodites, 5 days later, the 6-cm plates were chunked to ten 15 cm plates of high growth medium (HGM) covered with an *E. coli* lawn and grown six more days at 22°C. On the day of the harvest, worms were washed off the plates with M9 buffer, and pelleted at 3,200 g for 15 minutes several times (2-3) to reduce the volume and compact the worms into a 15 mL conical single pellet of approximate volume of 2 mL. The pellet was transferred to an aluminum foil sheet with a spatula, and the foil was dropped into liquid nitrogen to freeze the worm pellet. Frozen batches were stored frozen at-80°C until all the biological samples were collected. To prepare the frozen worm pellets for efficient lysis, the frozen pellets were pulverized using the Cell Crusher Kit and stored in 15 mL conical tubes at-80°C until the day of further processing.

### Sample lysis and desalting

To each frozen pulverized pellet, we added a TBS buffer with protease inhibitors (50mM Tris-HCl ph 8.0, 150 mM NaCl, Pierce HALT protease inhibitor cocktail) at 1:10 ratio and vortexed immediately to resuspend the pellet. Once resuspended, 20% SDS was added to a final concentration of 1%. The samples were quickly heated to 90°C by placing the lysate tubes in a boiling hot water bath and stirring continuously for 10 minutes. The samples were sonicated using short pulses at 40% amplitude for 1 min each, we did two sonication rounds. The samples were cooled down to room temperature and 8M urea was added to a final concentration of 2M. The samples were then centrifuged at 38,700 g for 30 min at 22°C. Supernatant (approximately 10 mL) was transferred to a new tube for further desalting step aimed to deplete endogenous biotin levels in the lysate using Zeba spin desalting columns (7 K molecular cutoff, 10mL each). The columns were equilibrated three times with 10 mL TBS buffer containing 2M urea by centrifugation at 1000 g for 10 min (or until the buffer was completely eluted from the resin). 10 mL of clarified sample was then loaded onto each equilibrated column and desalted by centrifugation at 1000 g for 10 min (or until the sample was completely eluted from resin) to remove free biotin.

### Depletion of His-tagged carboxylases

1 mL per sample of Pure Cube Ni-NTA agarose resin (Cube Biotech) was transferred to a 15 mL conical tube, centrifuged for 5 min at 1,000 g, and the supernatant was discarded. The Ni–NTA agarose was then equilibrated with 10 mL of Ni–NTA binding buffer (sodium phosphate buffer, 4M urea, pH 8.0) for 10 min, centrifuged for 5 min at 1,000 g, and the supernatant was discarded. The wash was repeated one more time and then mixed with the protein lysate with 3 volumes of Ni-NTA binding buffer in a 50 mL conical tube. The mix was incubated for 2 h at room temperature using a tube roller. The tube was centrifuged at 1,000 g for 5 min, and the supernatant was transferred to a new 50 mL conical tube with great care not to transfer any of the pelleted resin.

### Streptavidin magnetic bead isolation

Pierce Streptavidin magnetic beads (Thermo Fisher Scientific; 150 μL beads per sample) were washed three times with 1 mL of buffer 1 (50 mM Hepes–NaOH, pH 7.8, and 0.2% Tween-20), briefly incubated in the buffer, and then separated using magnetic separation to retain the beads while discarding the buffer. Washed beads were resuspended in a mix of 190 μL of buffer 1 and 10 μL of 100 mM Pierce Sulfo-NHS-Acetate (Thermo Fisher Scientific; dissolved in dimethyl sulfoxide), and incubated for 1 h at room temperature to acetylate free amines. Beads were washed three times with 1 mL of buffer 2 (50 mM ammonium bicarbonate and 0.2% Tween-20) to stop the reaction.

Carboxylase-depleted supernatant was mixed with the acetylated magnetic beads in a 50 mL conical tube and incubated overnight on a tube roller at room temperature. Beads were collected on a magnet rack. Unbound lysate was aspirated, and the beads were transferred to a 2 mL LoBind protein tube (Eppendorf) using 2% SDS wash buffer (150 mM NaCl, 1 mM EDTA, 2% SDS, 50 mM Tris–HCl, pH 7.4). Beads were washed twice with 2% SDS wash buffer, once with TBS-T buffer (150 mM NaCl, 50 mM Tris–HCl, 0.2% Tween-20, pH 7.6), twice with 1 M KCl-T wash buffer (1 M KCl, 1 mM EDTA, 50 mM Tris–HCl, 0.2% Tween-20, pH 7.4), twice with 0.1 M Na2CO3-T buffer (0.1 M Na2CO3, 0.2% Tween-20, pH 11.5), twice with 2 M urea-T buffer (2 M urea, 10 mM Tris–HCl, 0.2% Tween-20, pH 8.0) and five times with 50mM ammonium bicarbonate.

During each wash, 1 mL of buffer was used, beads were incubated on a rocking platform for 10 to 15 min and transferred to new tubes as often as possible. After the final wash, the pellet of beads was frozen at-80°C until further processing for mass spectrometry.

### On-beads digestion

0.2ug of trypsin in 20 ul 50mM NH4HCO3 was added to washed and ready to digest beads and incubated at 37°C for 4 hours. Another 0.2 ug of trypsin was added and incubated at 37°C overnight. The solution was separated from the magnetic beads and pH was adjusted to 3 with 10% formic acid. The sample was desalted with stage tip^72^ before LC-MSMS.

### Liquid Chromatography–Tandem Mass Spectrometry (LC***-***MS/MS)

Peptides were analyzed by nano-LC-MS/MS using a Vanquish Neo nanoLC system (Thermo Fisher Scientific) coupled to a timsTOF HT mass spectrometer (Bruker Daltonics). Samples were directly loaded onto a PepSep Ultra C18 column (75 µm × 25 cm, Bruker) and separated at a flow rate of 300 nL/min using a segmented linear gradient: 2–12% solvent B over 16 min, 12–20% B over 16 min, and 20–32% B over 12 min (solvent A: 0.2% formic acid in water; solvent B: 0.2% formic acid in acetonitrile). For DIA-PASEF acquisition, precursor ions with m/z values between 300 and 1200 were analyzed using eight DIA-PASEF scans per cycle, each consisting of three quadrupole isolation windows and 24 ion mobility steps spanning a range of 0.7–1.3 (1/K₀). Variable isolation window widths of 36–41 Th were applied for each ion mobility step. The ramp time and accumulation time were both set at 85 ms to make the ramp rate of 10.97 Hz.

Raw data were analyzed with predicted library combining UniProt reference proteome UP000001940 and Uniprot E.coli reference proteome UP000000625 as well as custom sequences in DIA-NN 2.02 Academia^73^ with following parameters: M-oxidation: no, cysteine-carbamidomethylation: no, N-term M excision: yes, min pep length set at 7, max pep-length set at 30. Mass range for MS1 set at 400-1200, protease was set as trypsin with max miss cleavage set at 1. The same settings were used for library-free search using DIA-NN 2.0.2 Academia with match between run selected.

## Data analysis

For the *C. elegans* data, differential abundance analysis was done using R package “promor”^74^. Spectral intensities were normalized using quantile normalization, missing values were imputed using the random forest algorithm, and linear statistical models for microarray data (limma) were applied to identify differentially abundant proteins. Mass spectrometry data have been deposited in the MassIVE repository (https://massive.ucsd.edu/ProteoSAFe/static/massive.jsp) and assigned the accession number MSV000101769)^75^. The dataset will be made public upon acceptance of the manuscript. A summary of the results is provided as supplemental data.

### Mammalian BioID and mass spectrometry analysis

BioID and mass spectrometry analyses were performed essentially as described.^76^ Briefly, stable HEK293 Flp-In T-REx cells were grown on 15-cm plates to approximately 75% confluency. Bait expression and proximity labeling were then induced simultaneously by addition of tetracycline (1 μg/ml) and biotin (50 μM) for 24 h. Cells were harvested in PBS, and biotinylated proteins were purified by streptavidin–agarose affinity chromatography. Proteins were digested on-bead with sequencing-grade trypsin in 50 mM ammonium bicarbonate (pH 8.5). Peptides were acidified by addition of formic acid to a final concentration of 2% (v/v) and dried by vacuum centrifugation. Dried peptides were resuspended in 5% (v/v) formic acid and analyzed on a TripleTOF 5600 mass spectrometer (SCIEX) coupled in-line to a nanoflow electrospray ion source and nano-HPLC system. Raw data were searched and analyzed within the ProHits LIMS^77^, and peptides were mapped to genes to determine prey spectral counts. High-confidence proximity interactions (SAINT score ≥ 0.80) were identified by SAINT analysis^27^ implemented within ProHits. Bait samples (biological duplicates) were compared against 8 independent negative control samples comprising 4 BirA-FLAG-only and 4 triple-FLAG-only expressing cell lines. Data have been deposited as a complete submission to the MassIVE repository (https://massive.ucsd.edu/ProteoSAFe/static/massive.jsp) and assigned the accession number MSV000101807. The ProteomeXchange accession is PXD078253. The dataset will be made public upon acceptance of the manuscript.

## Acknowledgements

We thank members of the Grant lab for critical comments and suggestions, Marie Mao for plasmid construction, Haiyan Zheng for expert mass spectrometry assistance, and Junjie Liu for expert microinjection. Instrumentation was supported by NIH grant 1S10OD036226 (TimsTOF HT and Vanquish NEO). This work was supported by a grant from the Polycystic Kidney Disease Foundation (pkdcure.org) to I.A.N., the Canadian Institutes of Health Research (CIHR) grant FDN 143301 to A-C.G., and NIH grant 5R01GM135326 to B.D.G. The Polycystic Kidney Disease Foundation had no role in study design, data collection and interpretation, or the decision to submit the work for publication.

## Abbreviation

ESCRT: Endosomal Sorting Complexes Required for Transport
MIG-14/Wls: Wntless
RME-8: Receptor Mediated Endocytosis-8
SCM-1/SCAMP: Secretory Carrier-Associated Membrane Protein
v-SNARE: vesicle-Soluble N-ethylmaleimide-sensitive factor Attachment protein Receptors
SNB-2: SyNaptoBrevin 2
VAMP: Vesicle-Associated Membrane Protein.

---

**Supplementary Fig. 1.**
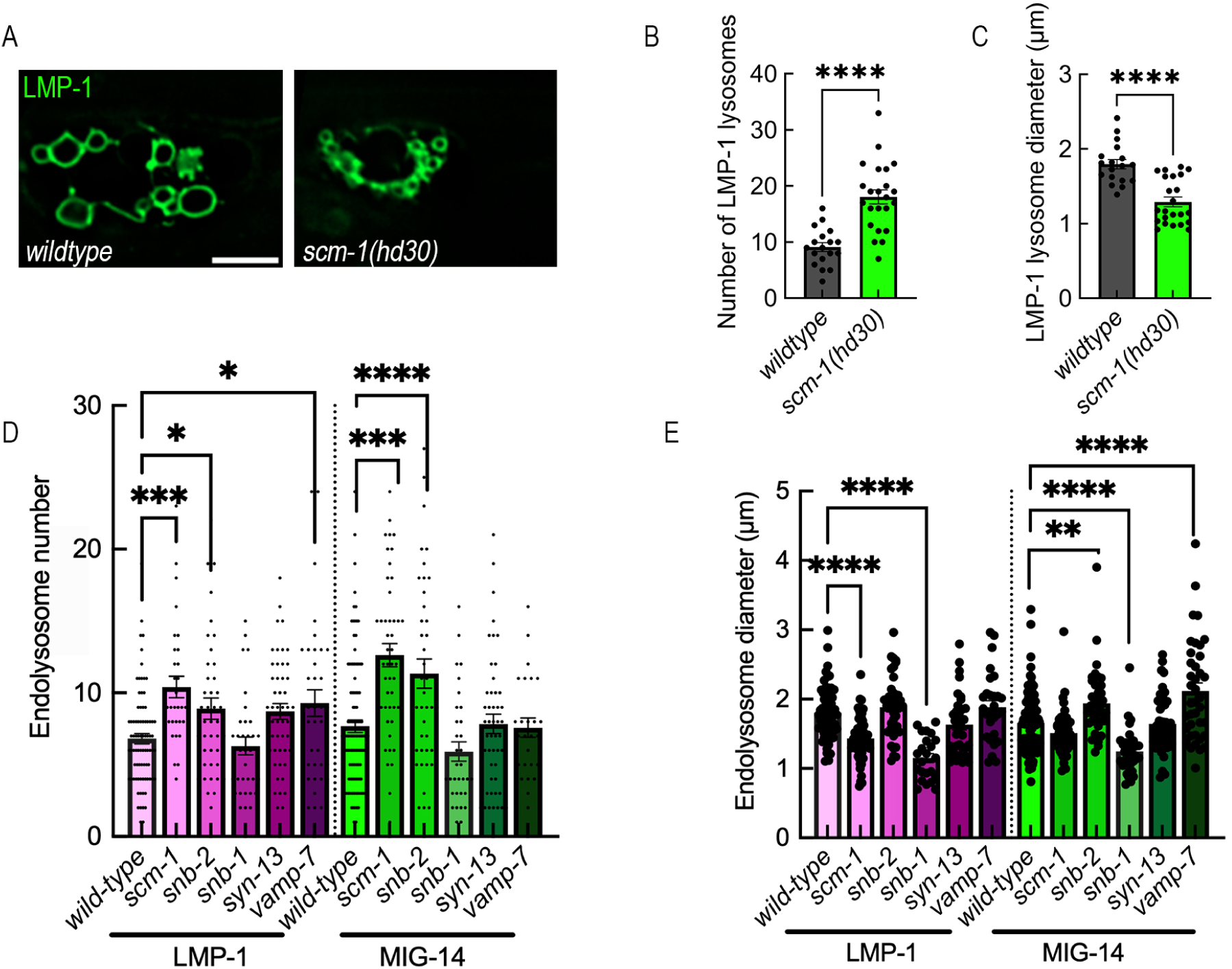
***snb-2* phenocopies *scm-1* endolysosomal number phenotypes** (A) LMP-1::GFP expressed in wildtype and *scm-1(hd30)* background (B-C) Quantification of LMP-1 vesicle number and diameter. *scm-1* mutants have smaller and more numerous lysosomes (D) Quantification of the number of MIG-14 and LMP-1 vesicles in control, *scm-1,* and SNARE mutant animals corresponding to images in Fig. 4G-L. *snb-2* mutants show an increase in lysosome number similar to *scm-1* mutants, whereas other SNARE mutants do not. (E) Quantification of diameter for MIG-14::GFP-positive vesicles and LMP-1::GFP labeled lysosomes in control, *scm-1,* and SNARE mutant animals corresponding to images in Fig. 4G-L. Data are mean± SEM; each data point represents average of one independent experiment (n>12 coelomocytes per experiment). no annotation indicated no signifigance, *p:5 0.05, **p:5 0.01, ***p:5 0.001 (one-way ANOVA) Scale bar is 5microns.

**Supplementary Fig. 2.**
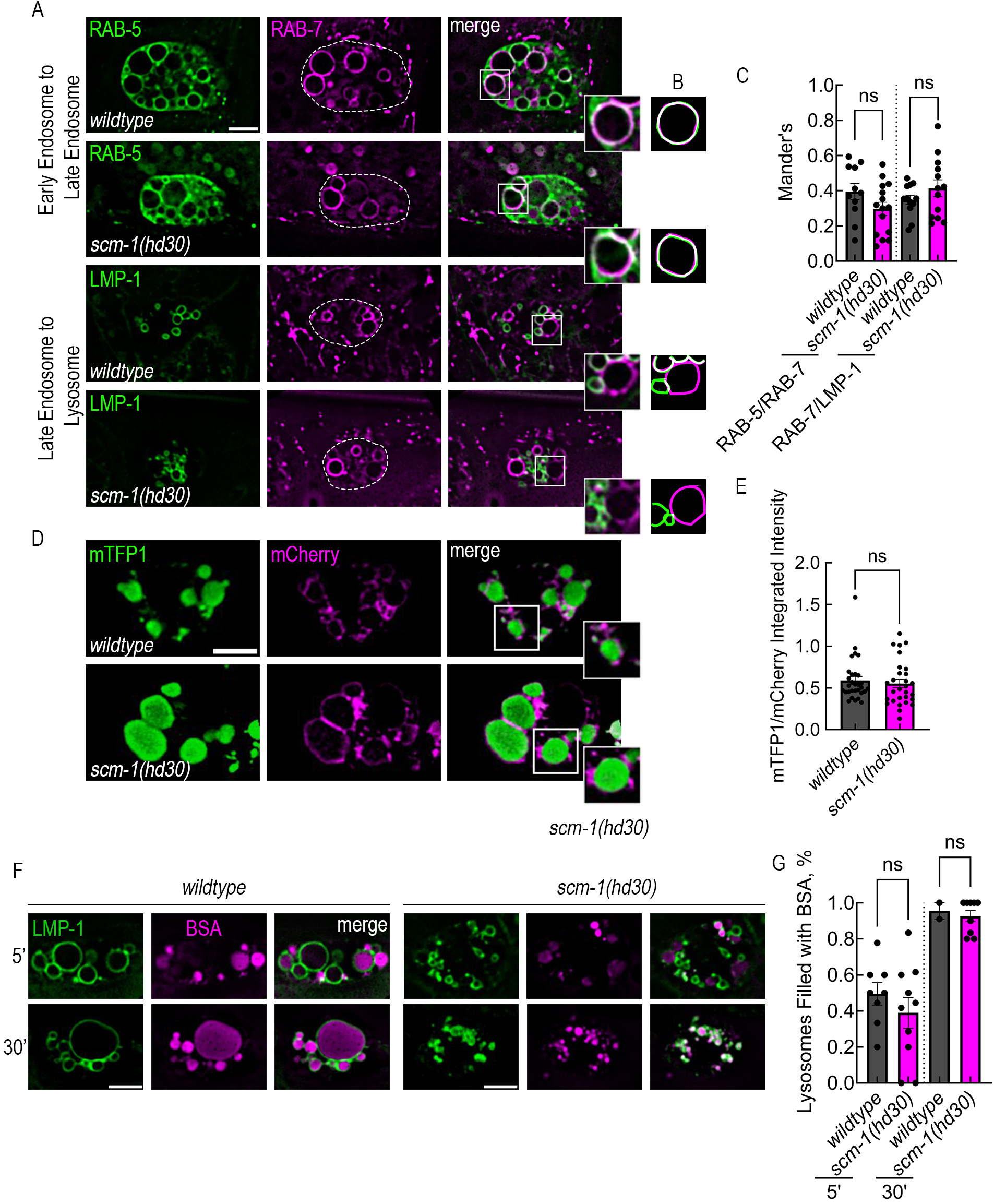
SCM-1 does not affect endosomal maturation, acidification, or fluid-phase transport. (A) The degree of colocalization of GFP::RAB-5 and endogenously tagged mScarlet::RAB-7, and the degree of colocalization of mScarlet::RAB-7 and LMP-1::GFP, is unchanged in *scm-1* mutants. Insets are 2-fold larger. (B) Cartoons of insets from A. (C) Quantification of colocalization by Manders’ coefficient. (D) Images of FIRE-pHly (mTFP1::LMP-1::mCherry) pH biosensor in coelomocytes. (E) mTFP1/mCherry ratio does not change in *scm-1* mutants. (F) Transport of endocytosed Cy5-BSA to lysosomes proceeds with normal kinetics in *scm-1* mutants. (G) Percent of lysosomes containing BSA at 5 and 30 minutes in wildtype and *scm-1* coelomocytes. Scale bar: 5 µm (A, D, F). Data are presented as mean± SEM. Each data point represents a single coelomocyte, data shown (n>9 for A,D and n>2 for F) is representative of three independent experiments.

**Table S1.** Strains used in this study.

| Strain name | Genotype | Reference |
| --- | --- | --- |
| AX7884 | *pod-2(syb1772[pod-2::His10]) II; mccc-1(syb1666[mccc-1::His10]) IV; pyc-1(syb1680[pyc-1::His10]) V; pcca-1(syb1626[pcca-1::His10]) X.* | <sup>71</sup> |
| RT3518 | *pwSi254[psnx-1::GFP::SCM-1(synthetic); pwls1022[psnx-1::AMAN-2::tagRFP]* | This study |
| RT3516 | *pwSi254[psnx-1::GFP::SCM-1(synthetic)]; pwls1053[psnx-1::tagRFP::RME-8]* | This study |
| RT3962 | *pwSi254[psnx-1::GFP::SCM-1(synthetic)]; pwls1038[psnx-1::tagRFP::HGRS-1]* | <sup>8</sup> |
| RT4878 | *pwSi544[psnx-1::mScarlet::SCM-1]; pwls50[plmp-1::LMP-1::GFP]* | This study |
| RT2800 | *pwls984[psnx-1::tagRFP::SNX-1]; pwls1039[psnx-1-citrine::HGRS-1]* | <sup>8</sup> |
| RT3482 | *pwls984[psnx-1::tagRFP::SNX-1]; pwls1039[psnx-1-citrine::HGRS-1]; scm-1(hd30)* | This study |
| RT2764 | *pwls1039[psnx-1::HGRS-1::citrine-unc54-3'UTR]* | <sup>8</sup> |
| RT3846 | *pwls1039[psnx-1::HGRS-1::citrine-unc54-3'UTR]; scm-1(hd30)* | This study |
| RT225 | *pwls94[psnx-1::GFP::SNX-1]* | <sup>13</sup> |
| RT4938 | *pwls94[psnx-1::GFP::SNX-1]; scm-1(hd30)* | This study |
| RT3864 | *pwls995[psnx-1::MIG-14::GFP-unc54 3'UTR]; pwls1054[psnx-1::tagRFP::RAB-5::unc54 3'UTR];* | This study |
| RT3865 | *pwls995[psnx-1::MIG-14::GFP-unc54 3'UTR]; pwls1054[psnx-1::tagRFP::RAB-5::unc54 3'UTR]; scm-1(hd30)* | This study |
| RT4922 | *pwls995[psnx-1::MIG-14::GFP]; xdKi59[wrmsScarlet::RAB-7]* | This study |
| RT4923 | *pwls995[psnx-1::MIG-14::GFP]; xdKi59[wrmsScarlet::RAB-7]; scm-1(hd30)* | This study |
| RT2732 | *pwls995[psnx-1::MIG-14::GFP]; cdls139(plmp-1::LMP-1::tagRFP]* | This study |
| RT4920 | *pwls995[psnx-1::MIG-14::GFP]; cdls139(plmp-1::LMP-1::tagRFP]; scm-1(hd30)* | This study |
| RT2702 | *pwls995[psnx-1::MIG-14::GFP]* | <sup>8</sup> |
| RT3849 | *pwls995[psnx-1::MIG-14::GFP]; scm-1(hd30)* | This study |
| RT258 | *pwls50[plmp-1::LMP-1::GFP + Cbr-unc-119(+)]; unc-119(ed3) II* | <sup>78</sup> |
| RT4772 | *pwls50[plmp-1::LMP-1::GFP + Cbr-unc-119(+)]; scm-1(hd30)* | This study |
| GS1826 | *arls36[hsp::ssGFP]* | <sup>79</sup> |
| RT4939 | *arls36[hsp::ssGFP]; scm-1(hd30)* | This study |
| RT688 | *pwls281 [ppie-1::CAV-1::GFP]* | <sup>50</sup> |
| RT4891 | *pwls281[ppie-1::CAV-1::GFP]; scm-1(hd30)* | This study |
| RT4876 | *xdKi59[wormScarlet::RAB-7]; pwls439[prab-5::GFP::RAB-5(genomic)]* | This study |
| RT4877 | *xdKi59[wormScarlet::RAB-7]; pwls439[prab-5::GFP::RAB-5(genomic)]; scm-1(hd30)* | This study |
| RT4940 | *xdKi59[wormScarlet::RAB-7]; pwls50 [plmp-1::LMP-1::GFP]* | This study |
| RT4941 | *xdKi59[wormScarlet::RAB-7]; pwls50 [plmp-1::LMP-1::GFP]; scm-1(hd30)* | This study |
| RT4683 | *pwSi603[pcup4::FirepHLY ::let858::3'UTR]* | This study |
| RT4943 | *pwSi603[pcup4::FirepHLY ::let858::3'UTR]; scm-1(hd30)* | This study |
| RT4944 | *pwSi589[psnx-1::GFP::SNB-2]; pwSi544[psnx-1::mScarlet::SCM-1]* | This study |
| RT4953 | *pwSi598[psnx-1::GFP::SNB-1]; pwSi544[psnx-1::mScarlet::scm-1]* | This study |
| RT4954 | *pwSi597[psnx-1::GFP::SYX-7]; pwSi544[psnx-1::mScarlet::SCM-1]* | This study |
| RT4955 | *pwSi586[psnx-1::GFP::VAMP-7]; pwSi544[psnx-1::mScarlet::SCM-1]* | This study |
| RT4886 | *pwls995[psnx-1::MIG-14::GFP]; cdl139(plmp-1::LMP-1::tagRFP); snb-2(tm3725)* | This study |
| RT4888 | *pwls995[psnx-1::MIG-14::GFP]; cdl139(plmp-1::LMP-1::tagRFP); snb-1(md247)* | This study |
| RT4887 | *pwls995[psnx-1::MIG-14::GFP]; cdl139(plmp-1::LMP-1::tagRFP); syx-7(tm1764)* | This study |
| RT4921 | *pwls995[psnx-1::MIG-14::GFP]; cdl139(plmp-1::LMP-1::tagRFP); vamp-7(tm6588)* | This study |
| RT4956 | *pwSi589[psnx-1::GFP::SNB-2 let858]; pwSi577[psnx-1::LMP-1::mKate2]* | This study |
| RT4957 | *pwSi589[psnx-1::GFP::SNB-2]; pwSi577[psnx-1::LMP-1::mKate2]; scm-1(hd30)* | This study |

## Notes

### Competing Interest Statement

The authors have declared no competing interest.

